# Co-administration of macropinocytosis inhibitory nanoparticles (MiNP) for enhanced nanoparticle circulation time and target tissue accumulation following subcutaneous injection

**DOI:** 10.1101/2020.08.26.267054

**Authors:** Trevor Stack, Yu-gang Liu, Molly Frey, Sharan Bobbala, Michael Vincent, Evan Scott

## Abstract

A signficant barrier to the application of nanoparticles for precision medicine is the mononuclear phagocyte system (MPS), a diverse population of phagocytic cells primarily located within the liver, spleen and lymph nodes. The majority of nanoparticles are indiscriminantly cleared by the MPS via macropinocytosis before reaching their intended targets, resulting in side effects and decreased efficacy. This work demonstrates that the biodistribution and desired tissue accumulation of targeted nanoparticles can be significantly enhanced by co-injection with polymeric micelles containing the actin depolymerizing agent **latrunculin A**. These macropinocytosis inhibitory nanoparticles (MiNP) were found to selectively inhibit non-specific uptake of a second “effector” nanoparticle *in vitro* without impeding receptor-mediated endocytosis. In tumor bearing mice, co-injection with MiNP in a single multi-nanoparticle formulation significantly increased the accumulation of folate-receptor targeted nanoparticles within tumors. Furthermore, subcutaneous co-administration with MiNP allowed effector nanoparticles to achieve serum levels that rivaled a standard intravenous injection. This effect was only observed if the effector nanoparticles were injected within 24 h following MiNP administration, indicating a temporary avoidance of MPS cells. Co-injection with MiNP therefore allows reversible evasion of the MPS for targeted nanoparticles and presents a previously unexplored method of modulating and improving nanoparticle biodstribution following subcutaneous administration.

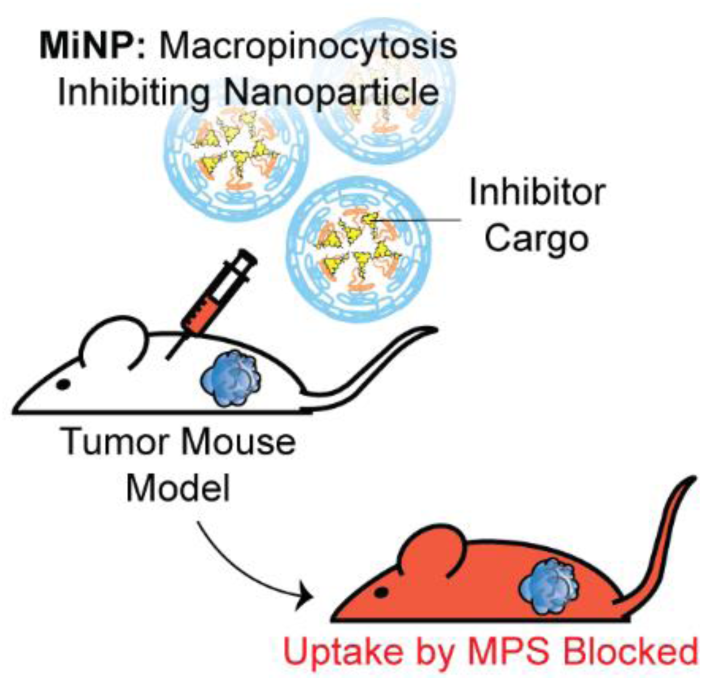

TOC Text: Polymeric macropinocytosis inhibiting nanoparticles reduce non-specific uptake of an “effector” nanoparticle by cells of the mononuclear phagocyte system. This macropinocytosis specific inhibition allows for greater accumulation and uptake of targeted nanoparticles in tissues of interest thereby increasing their efficacy and reducing side effects.

## Introduction

Nanoparticles are versatile carriers that can improve and often specify the stability, circulation time, and biodistribution of therapeutic molecules^[1]^. Despite these advantages, rapid clearance of nanoparticles by the mononuclear phagocyte system (MPS)^[2]^ remains a significant barrier to their applications in precision medicine. The MPS consists of circulating and organ-resident phagocytic cells, which internalize nanoparticles and eventually clear them through the liver^[3–6]^. A comprehensive survey of the literature reported that a median average of only 0.7% of administered nanoparticles successfully reach solid tumors despite the use of surface-conjugated targeting moieties like antibodies, peptides or aptamers^[7]^. Clearance by MPS cells occurs primarily in the liver, spleen and lymph nodes through a number of endocytic pathways including clathrin-mediated and clathrin-independent pathways, macropinocytosis, and phagocytosis^[8]^. If these pathways are temporarily inhibited in MPS cells prior to or in conjunction with the introduction of therapeutic nanoparticles, the bioavailability and therapeutic efficacy of these nanoparticles would likely increase significantly^[9]^. Developing nanoparticles with “stealth” properties to avoid this non-specific uptake remains a critical objective for nanomedicine and many different strategies such as PEGylation and CD47 “don’t eat me” peptides, have been tried with variable levels of effectiveness^[10–12]^. Combinatorial strategies employing multiple different stealth strategies are needed to further reduce clearance by the MPS and increase nanomaterial utility *in vivo*.

We have previously demonstrated that diverse nanoparticle morphologies can be self-assembled from poly(ethylene glycol)-*block*-poly(propylene sulfide) (PEG-*b*-PPS) copolymers to function as customizable and non-toxic drug delivery vehicles^[13,14]^. The nanostructure morphology and route of administration dictate the biodistribution of PEG-*b*-PPS nanoparticles, allowing the passive and preferential targeting of different phagocytic cell populations *in vivo* without the need for surface-conjugated targeting ligands^[13,15–18]^. Of note, spherical solid core PEG-*b*-PPS micelles are primarily taken up by liver macrophages following intravenous (IV) administration^[16]^ and monocyte populations in draining lymph nodes and spleen following subcutaneous (SC) administration^[15]^, both of which are key components of the MPS^[19]^. These micelles have also been previously shown to reduce the cytotoxicity of small molecule drugs such as celastrol.^[20]^ Here, we employ macropinocytosis inhibitory nanoparticles (MiNP) to reduce non-specific nanoparticle uptake by the MPS and enhance their accumulation within target tissues. MiNP are comprised of PEG-*b*-PPS micelles containing Latrunculin A (LatA), a well-known and transient actin depolymerizing agent^[21]^. LatA is most commonly used to temporarily inhibit macropinocytosis^[22]^ by phagocytic cells during *in vitro* assays to investigate mechanisms of cell endocytosis. Furthermore, LatA is hydrophobic and thus not amenable to direct administration via SC or IV routes. We have previously shown that LatA retains its inhibitory effects by disrupting the cell cytoskeleton when encapsulated in PEG-*b-*PPS micelles but without toxicity^[23,24]^.

We selected to investigate and optimize a co-administration strategy wherein MiNP are injected before and/or simultaneously with a model “effector” nanoparticle, which represents a nanoparticle employed for either diagnostic or therapeutic applications that will be enhanced by decreased MPS clearance. For the purposes of this proof of concept study, the employed model effector nanoparticle is a fluorescent PEG-*b*-PPS micelle (E-MC). The inhibitory effects of MiNP were characterized *in vitro* using macrophages and *in vivo* in a B16F10 melanoma tumor bearing mouse model. Furthermore, E-MC with surface-decorated folate (E-MC(FA)) were used to explore the ability of MiNP to enhance the accumulation of a targeted E-MC within folate receptor-expressing solid tumors (**Figure 1**). We find that IV or SC injection of MiNP temporarily inhibit the non-specific MPS uptake of a subsequent chasing dose of an E-MC by increasing blood concentration and tumor accumulation compared to E-MC administered alone. Of note, this MiNP co-administration strategy significantly improved the SC injection of chase effector nanoparticles, achieving bioavailability of E-MC on par with IV injections.

**Figure 1:**
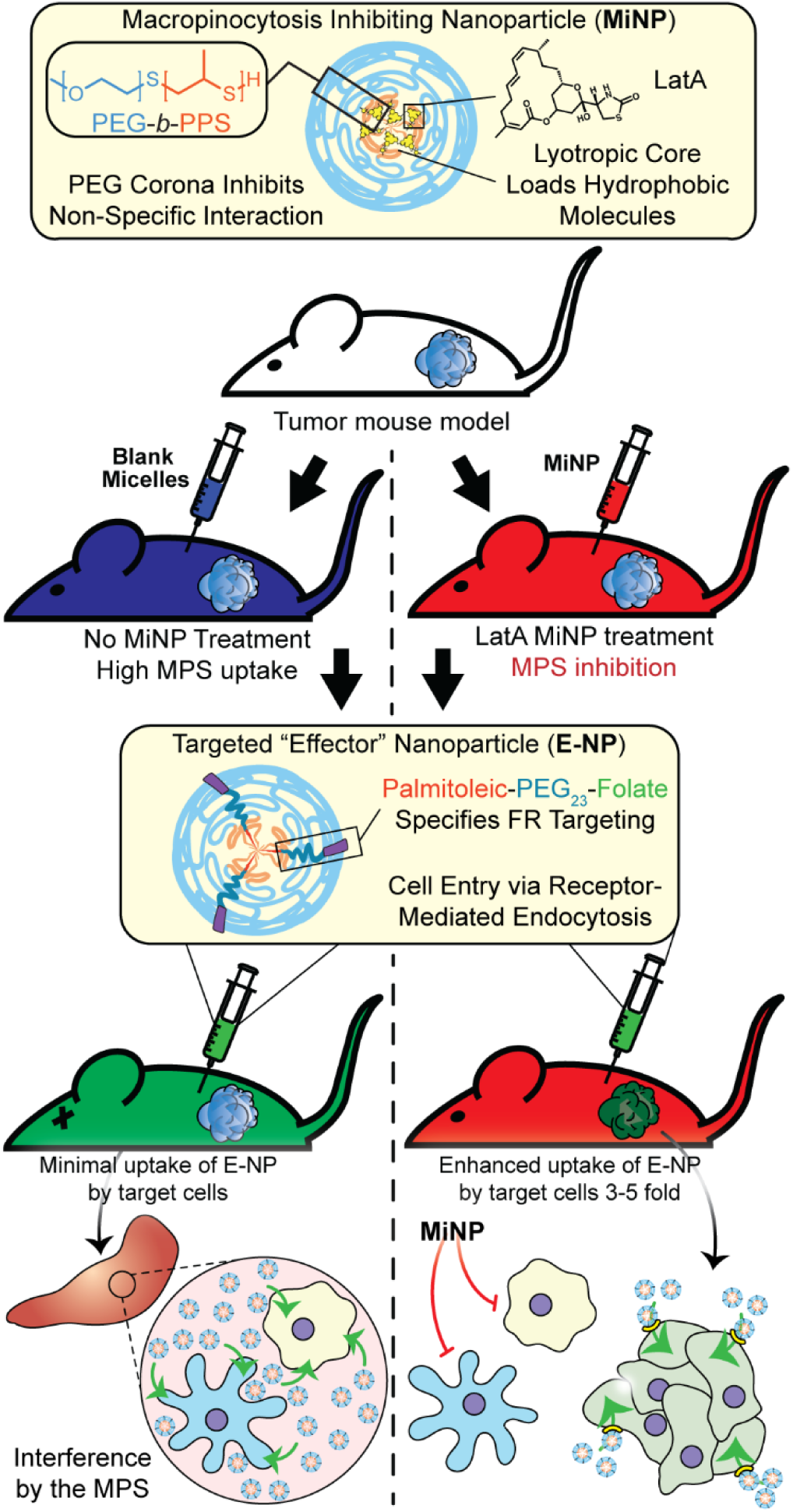
Schematic of tumor bearing mice being co-administered with latrunculin A-loaded macropinocytosis inhibiting nanoparticles (MiNP). MiNP were developed and evaluated for their effect on the accumulation of a targeted “effector” nanoparticle via subcutaneous and intravenous injection. As MiNP interferes with macropinocytosis but not receptor-mediated endocytosis, pre-and/or co-injection of MiNP with an effector nanoparticle displaying targeting ligands allows enhanced uptake by cells expressing the target receptor. As an example, MiNP are shown enhancing the targeting of receptors highly expressed within tumor microenvironments by interfering with off-target MPS clearance.

## Results

LatA-loaded ((+)MiNP) and unloaded controls ((-)MiNP) were self-assembled from PEG_45_-*b*-PPS_23_ using the co-solvent evaporation method^[25]^. Dynamic light scattering (DLS) was used to determine the z-average and polydispersity of the different formulations (**Table S1**). Confirmation of the micelle structure was obtained using cryogenic transmission electron microscopy and small angle X-ray scattering (SAXS) studies (**Figure 2a, b**). SAXS Scattering curves of both (+)MiNP and (-)MiNP were successfully fitted with a micelle model indicating retention of micellar nanostructures for (+)MiNP after loading with LatA (Figure 2b). The core radius and approximate diameter of both (-)MiNP and (+)MiNP obtained using SAXS model fits are reported in **Table S2**. LatA quantification and loading within the micelles was determined by HPLC-UV as previously reported^[24]^ and allowed for all formulations to be referenced based on their LatA content (Table S1). These data are consistent with our previous findings that encapsulation of LatA by PEG-*b*-PPS micelles does not alter their physical structure or polydispersity.

**Figure 2:**
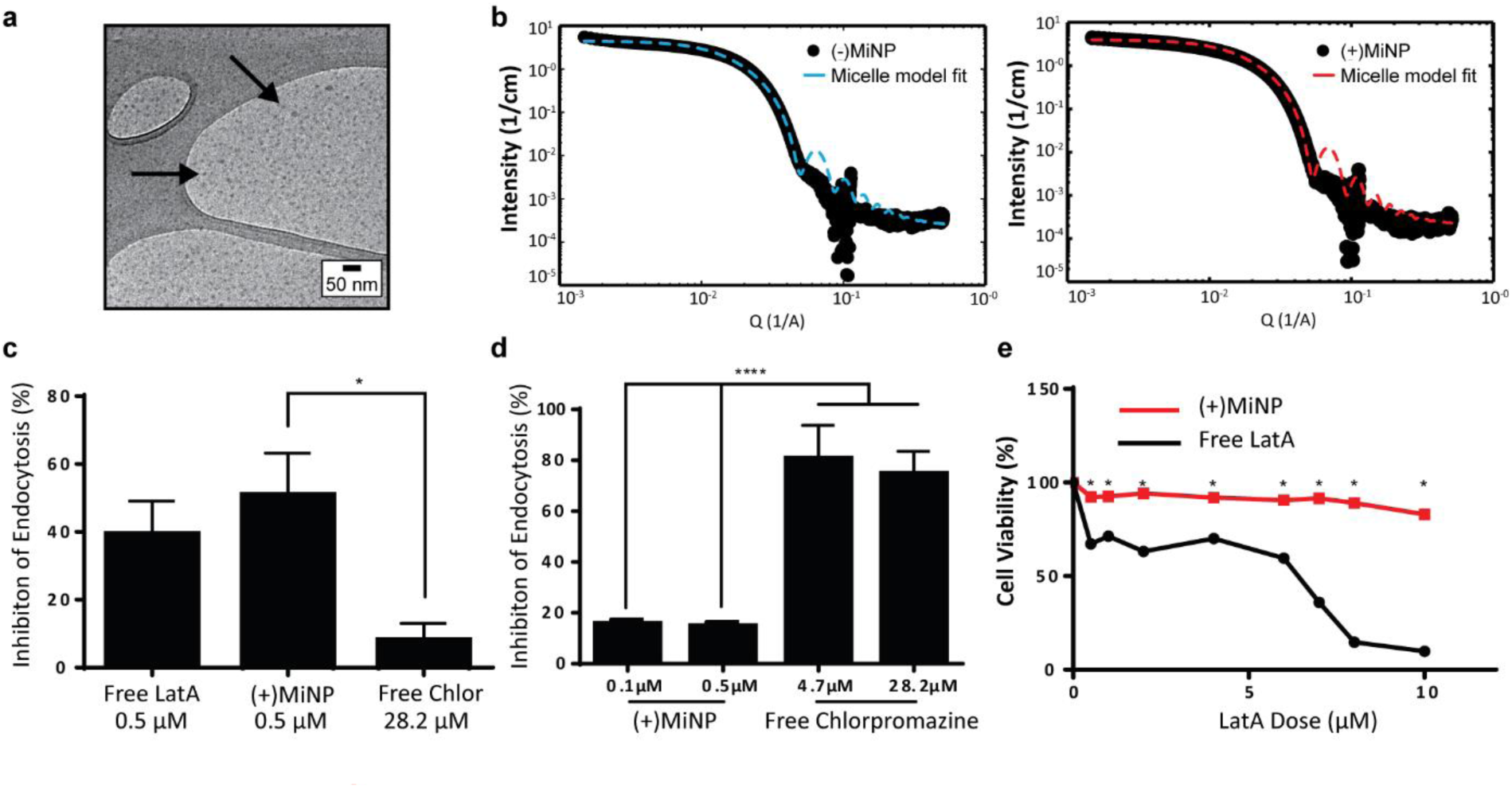
LatA retains its endocytic inhibition properties and does not change the size of PEG-*b*-PPS micelles when encapsulated. a) Cryogenic transmission electron microscopy (CryoTEM) of MiNP visually confirms retention of micellar structures. b) MiNP with ((+)MiNP) and without ((-)MiNP) loaded LatA were characterized via small angle x-ray scattering (SAXS) and fitted with a micelle model fit using SASView. c) Free LatA and (+)MiNP significantly inhibited non-receptor mediated endocytosis by RAW264.7 macrophages as compared to clathrin-mediated endocytosis inhibitor chlorpromazine. Cells were treated with each inhibitor for 2 h followed by 30 min of incubation with pHrodo dextran prior to analysis by flow cytometry. Data are shown as percent inhibition of endocytosis normalized to untreated macrophages. N=3 *p*<0.001. d) In comparison, uptake of transferrin conjugated pHrodo dextran by macrophages via receptor-mediated endocytosis was significantly inhibited by chlorpromazine compared to (+)MiNP. N=3 *p*<0.001. e) Loading within (+)MiNP significantly decreased the toxicity of LatA. Macrophages were incubated with various doses of free LatA or (+)MiNP for 4 h and assessed by flow cytometry for viability via the Zombie Aqua live/dead assay. N=3 *p*<0.05.

To investigate macropinocytosis inhibition by MiNP and determine whether this inhibition still permits uptake via receptor-mediated endocytosis, (+)MiNP were compared to chlorpromazine, an inhibitor of receptor-mediated endocytosis. These distinct mechanisms of endocytosis were evaluated using dextran conjugated pHrodo dye and transferrin conjugated pHrodo dye to respectively quantify effects of (+)MiNP and chlorpromazine on macropinocytosis (dextran) and receptor-mediated endocytosis (transferrin). Free LatA, (+)MiNP, and free chlorpromazine were incubated with RAW264.7 macrophages for 2 hours and subsequently washed and then chased with dextran-pHrodo (Figure 2c) or transferrin-pHrodo (Figure 2d). After 30 minutes of incubation, the cells were harvested and analyzed by flow cytometry to quantify and compare uptake of dextran-pHrodo via macropinocytosis and transferrin-pHrodo by transferrin-receptors. In the case of dextran (macropinocytosis), (+)MiNP and free LatA both showed much stronger inhibition of uptake than free chlorpromazine (Figure 2c). In the case of transferrin (receptor mediated endocytosis), free chlorpromazine at both high and low doses significantly inhibited endocytosis compared to (+)MiNP, which had minimal impact (Figure 2d). These results demonstrate that the functional aspect of LatA is not significantly altered by encapsulation in PEG-*b*-PPS micelles. Importantly, MiNP did not impede uptake of transferrin via transferrin receptors, suggesting that MiNP could be employed in a multi-nanoparticle strategy to inhibit non-specific uptake of a targeted chase nanoparticle while simultaneously permitting receptor-mediated targeting of specific cell populations. As LatA has been shown to be cytotoxic at higher doses and has been used as a cytotoxic agent^[26,27]^, we sought to evaluate the cytotoxicity of MiNP on our target cell population of macrophages. After 2 hours of exposure to various doses of (+)MiNP and Free LatA, (+)MiNP treated macrophages remained highly viable at all tested concentrations, while free LatA treated macrophages demonstrated significant toxicity at doses of 0.5 μM and above (Figure 2e). This is consistent with our previous findings in which encapsulation of small molecule drug Celastrol reduced its cytotoxicity in vitro^[20]^.

Having characterized MiNP in vitro, we next investigated different *in vivo* dosing regimens to evaluate the effect of MiNP on the uptake and biodistribution of a subsequently injected blank effector nanoparticle (E-MC) without targeting moieties (**Figure 3a**). Importantly, we sought to determine whether (+)MiNP could be administered SC with the E-MC in the same formulation versus administered as a separate injection prior to administration of the E-MC. For quantification of cell uptake, E-MC were labelled with DiR. We have previously demonstrated that Vybrant lipophilic dyes are stably retained within PEG-*b*-PPS nanocarriers for *in vivo* applications and flow cytometric analysis^[14,28,29]^. First, a pre-injection strategy was tested where (+/-)MiNP were administered twice at both 24 h and 4 h prior to injection of the chase nanoparticle. As we have previously published that the height of PEG-*b*-PPS micelle uptake by the MPS occurs at the 24 h timepoint^[16]^, we suspected that this regime would extensively pre-condition and shut down the MPS to avoid non-specific clearance of the E-MC. A simplified procedure was also evaluated wherein the (+/-)MiNP were administered alone just once to pre-condition the mouse, which was then followed 24 h later by a co-injected dose of a (+/-)MiNP and E-MC multi-nanoparticle formulation (**Figure 3a**). In both regimens, the same total micelle dosage of (+)MiNP or (-)MiNP and E-MC were administered. The total dosage of (-)MiNP and E-MC administered was equal to a LatA dose of 100 μL of 7 μM (+)MiNP solution (approx. 0.55 mg/kg). This dosage was based upon previously reported intraperitoneal treatments of mice^[26]^ and our *in vitro* toxicity assessment in RAW264.7 macrophages (Figure 2e). Mice were sacrificed 24 h after the chase injection, and organs were harvested for analysis by flow cytometry. Dendritic cells (DCs) and macrophages, the two key phagocytes of the MPS, were identified via antibody staining and the amount of E-MC in each cell population was quantified (**Figure S1**). In the spleen, (+)MiNP treatment with both regimes showed significantly less E-MC uptake by MPS cells than mice injected with (-)MiNP (Figure 3b). In the liver, the multi-nanoparticle co-injection (+)MiNP/E-MC formulation had significantly less chase particle uptake than (-)MiNP/E-MC in all cell types, while the 4 h pre-injection regimen did not. These results verify that a MiNP strategy indeed inhibits uptake of a second effector nanoparticle in both the spleen and liver following SC injection. Furthermore, this indicates that in both organs, MPS phagocytes are affected by the multi-nanoparticle (+)MiNP/E-MC co-injection dosing method at a greater or equivalent level than the 4 h separate pre-injection method. Additionally, the co-injection method is simpler to administer and would be preferred for any future translation of this system. As such, the co-injection multi-nanoparticle method was deemed superior and was the method of choice for future experiments.

**Figure 3.**
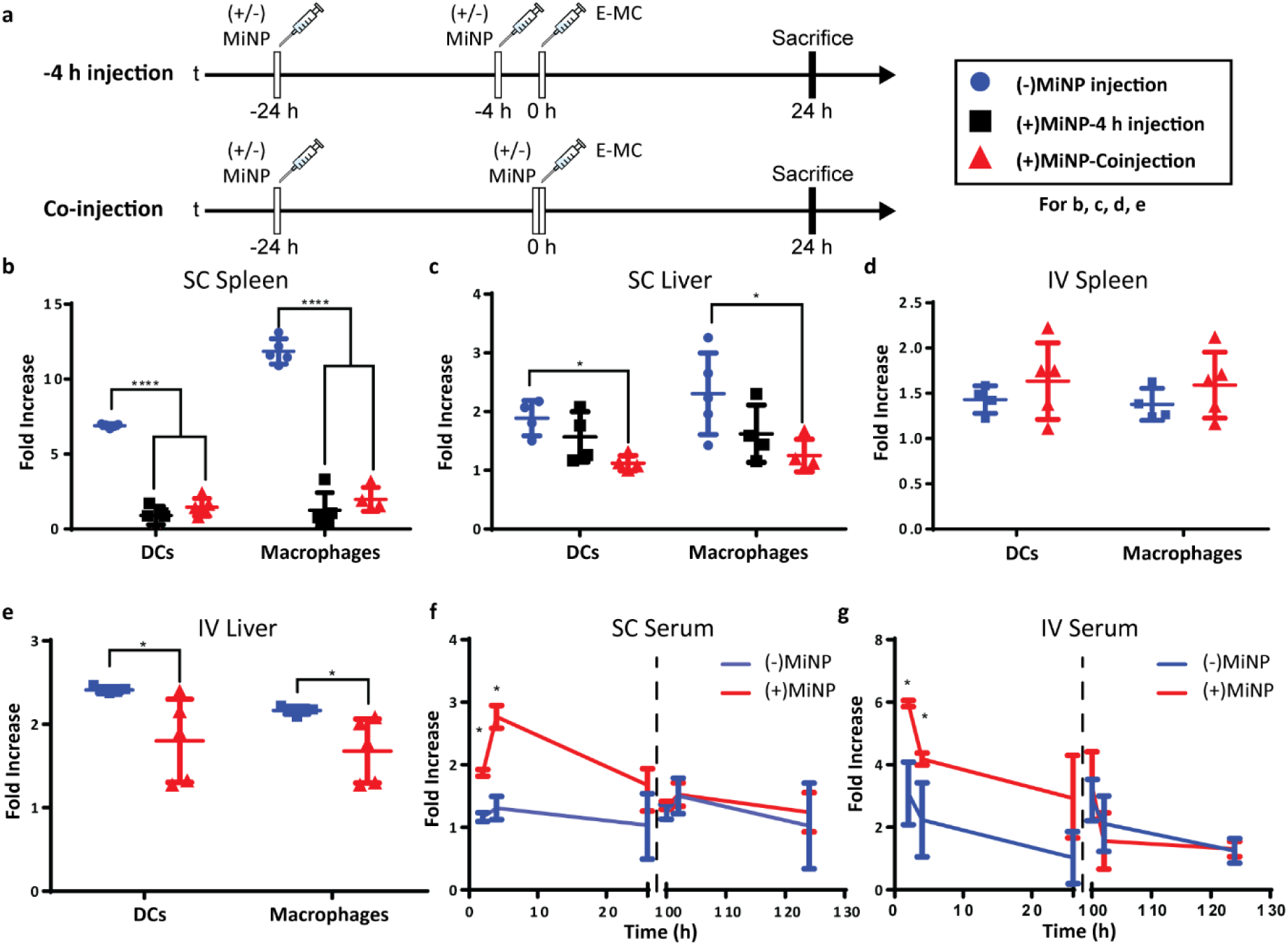
Latrunculin A loaded MC ((+)MiNP) co-injection and -4 h pre-injection lead to similar effector particle biodistributions. a) Timeline showing the injection times for the co-injection and −4 h injection methods. (+/-)MiNP indicated an injection of either (+)MiNP or (-)MiNP, and effector micelle injections are indicated by E-MC. All mice were sacrificed at 24 h post E-MC injection. Comparison of the two different injection strategies for (b, c) subcutaneous (SC) and (d, e) intravenous routes (IV) of injection. In all cases, mice were injected subcutaneously with 100 μL 7μM LatA (+)MiNP or (-)MiNP and E-MC were labelled with DiR for flow cytometric quantification of cellular uptake within the spleen and liver. Data are reported as fold increase median fluorescence intensity of the E-MC over an untreated control. N=5 *p*<0.0001. To assess the transience of the MiNP effect, mice were injected SC (f) or IV (g) with (+/-)MiNP and E-MC according to the co-injection method, and serum levels of E-MC were evaluated by fluorescence spectroscopy. Mice were then rested for 72 hours and injected again with only E-MC to determine whether the inhibitory effect remained. N=3 for 2 h and 4 h timepoints and N=6 for 24 h timepoints, \**p*<0.05.

We next sought to compare the effects of MiNP following SC versus IV administration. A majority of nanotherapeutics are administered IV out of necessity, as nanoparticles are rapidly cleared by phagocytes during lymphatic drainage. As IV administration must be performed by healthcare professionals, enhancing SC administration to achieve IV-level biodistribution of nanoparticles would permit unskilled administration and more flexible dose schedules, possibly increasing patient compliance and access to treatment. The co-injection protocol from SC injections was therefore followed for IV injections and the dose remained consistent at 100 μL of 7 μM LatA (+)MiNP. IV injection had a distinctly different uptake profile than SC injections, demonstrating no difference in E-MC uptake in the spleen (Figure 3d). However, in the liver, treatment with (+)MiNP decreased uptake of E-MC in DC and macrophage populations, indicating altered biodistribution within two MPS cell types of interest (Figure 3e).

We next evaluated serum levels of chase nanoparticles after SC and IV administration of (+)MiNP. LatA inhibits actin polymerization by binding actin at a 1:1 ratio and consuming intracellular LatA^[30]^. Thus, its effects should decrease over time due to continuous LatA depletion without replenishment. We therefore investigated the transient effects of (+)MiNP by evaluating whether the inhibitor’s effects would diminish within 100 h of the initial injection. Mice were divided into a (+) MiNP group that was administered a (+)MiNP/E-MC co-injection and a (-)MiNP control group that was administered (-)MiNP/E-MC. Blood (100 μL) was collected from each mouse at 2 h, 4 h and 26 h post SC or IV co-injection, and serum was isolated and analysed to assess E-MC content by spectrophotometry. Following final blood collection at 26 h, mice were rested for 74 h and subsequently injected with E-MC a second time at 100 h to assess any residual effects of the original MiNP administration. Blood (100 μL) was again collected from mice at 102 h, 104 h, and 126 h relative to the initial MiNP administration to quantify E-MC content by spectrophotometry. (+)MiNP treatment increased E-MC content in serum at 2 h and 4 h post injection following both IV and SC administration. The SC injections resulted in a delay in reaching the maximum serum level of the chase, which occurred at 4 h and is indicative of the time required for the nanoparticles to drain from the SC tissue and reach systemic circulation. After the mice were rested, there was no difference in the E-MC serum content between mice administered (+)MiNP or (-)MiNP after the second chase injection (Figure 3f, g). This indicates the mice returned to a baseline processing of E-MC by 100 h post (+)MiNP treatment for both routes of administration. Another interesting observation from this data was that the (+)MiNP SC injection group had a similar E-MC serum content at 4 h as the (-)MiNP IV injection group, suggesting that (+)MiNP administered subcutaneously are able to achieve a similar amount of E-MC serum concentration as the typically used IV injection ((-)MiNP treatment). This effect in conjunction with a therapeutic payload would allow access to a host of different dosing strategies for existing nanotherapeutics as well as easier administration.

Having shown in vitro that (+)MiNP can trasiently inhibit macropinocytosis while still allowing receptor-mediated endocytosis, we next sought to investigate whether (+)MiNP could enhance the uptake of chase nanoparticles targeting a specific cell receptor *in vivo*. The well-established B16F10 melanoma mouse model was chosen to compare the targeting of intratumoral folate receptors following IV and. SC routes of administration. B16F10 mouse melanoma cells have increased expression of folate receptors and folate decorated nanoparticles have been used by other groups to successfully target these cells^[31]^. We therefore synthesized a [folate] – [PEG linker] – [palmitoleic acid lipid anchor] (FA-PEG-PA) amphiphilic construct for stable incorporation into self-assembled PEG-b-PPS micelles (Figure 1). Briefly, folate was attached to a PEG1k-amine spacer that was then linked to a palmitoleic acid tail using EDC chemistry. The resulting FA-PEG-PA construct was incorporated into micelles by shaking overnight in phosphate buffered saline, allowing the palmitoleic acid anchor to partition into the hydrophobic PPS core of the micelle. We have previously demonstrated that such lipid anchored constructs could be stably retained within self-assembled PEG-*b*-PPS nanoparticles for controlled surface display of targeting moieties, such as peptides^[32]^. The formation of folate displaying micelles (E-MC(FA)) at controllable molar ratios of PEG-*b-*PPS polymer to FA-PEG-PA construct was confirmed using UV-Vis spectroscopy (**Figure S2**). An initial *in vitro* assessment by flow cytometry confirmed a significantly higher uptake of E-MC(FA) compared to E-MC following incubation with B16F10 melanoma cells for 1 h (**Figure S3**).

To investigate the enhancement in targeted delivery of MiNP co-administration, we compared the effect of our strategy on the uptake of folic-acid targeted E-MC(FA) vs. non-targeted E-MC. Mice were first inoculated with B16F10 melanoma cells, which were allowed to grow for approximately two weeks. Following an initial (+)MiNP or (-)MiNP injection at the −24 h timepoint, E-MC(FA) and E-MC were then coinjected with (+)MiNP or (-)MiNP using either SC or IV routes of administration (**Figure 4a**). E-MC(FA) and E-MC uptake was assessed in the liver, lymph nodes, spleen and tumor using flow cytometry. Vybrant DiR was used to identify both E-MC and E-MC(FA). IV injection of (+)MiNP increased E-MC uptake in tumor and lymph node CD45-cells(non-immune) as well as lymph node macrophages (**Figure 4b-d**). SC injection of (+)MiNP increased E-MC content in serum at 24 h as well as in liver and splenic DCs and macrophages (**Figure S4, Figure 4e-g**). No differences were found in any of the tumor cell subsets using SC administration. These data indicate that IV injection of MiNPs could facilitate increased tumor targeting of an effector therapeutic, but that SC would not.

**Figure 4.**
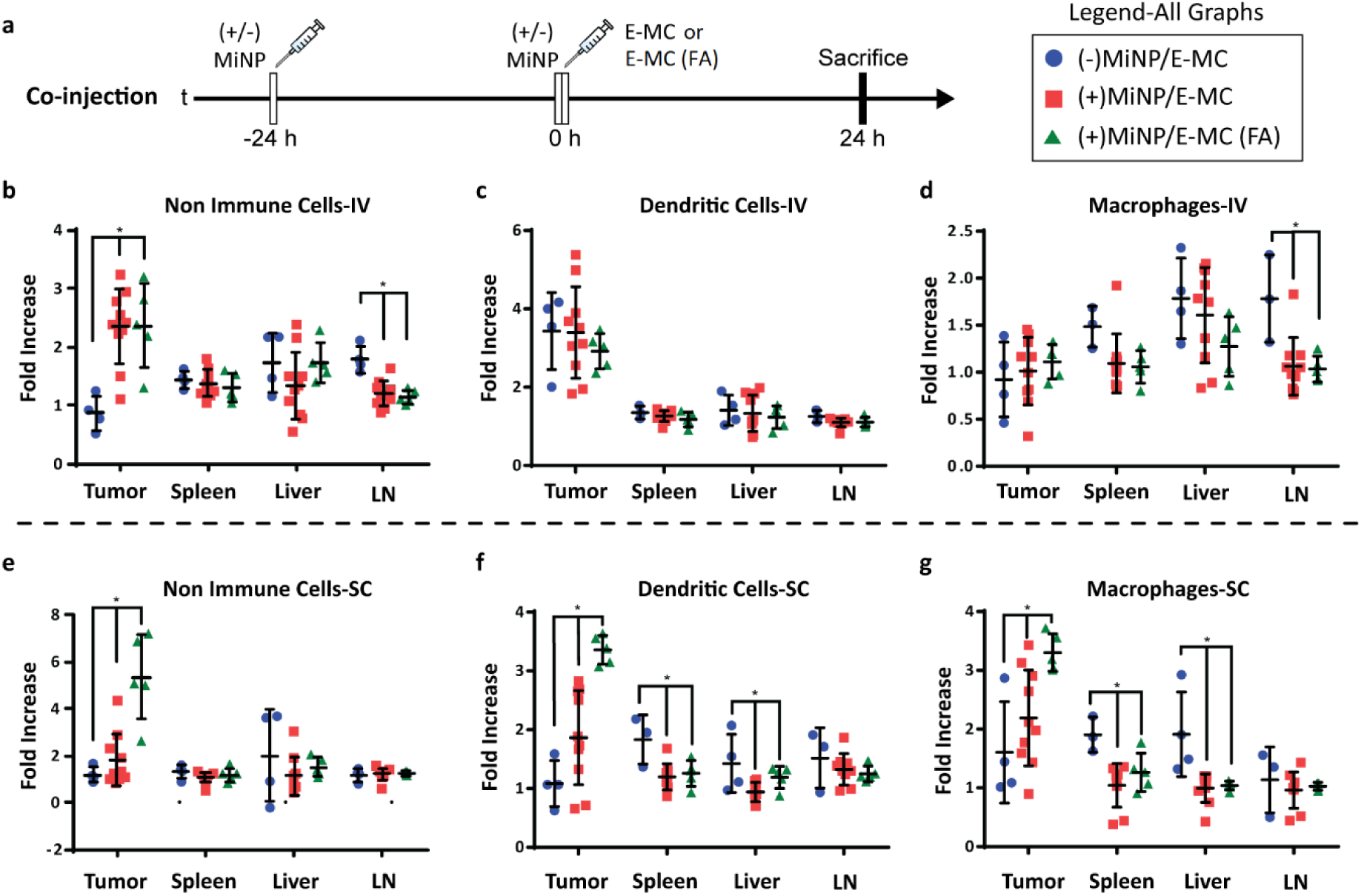
(+)MiNP treatment increases the accumulation of folate-targeted E-MC (E-MC(FA)) in B16F10 tumors following SC injection. a) Timeline of injection protocol assessing the tumor-targeting co-injection method. (+/-)MiNP indicates an injection of either (+)MiNP or (-)MiNP. Mice were sacrificed 24 h after the co-injection for analysis by flow cytometry. Results are shown for IV (b, c, d) and SC (e, f, g) injections of 3 co-injection modalities: (-)MiNP treatment/E-MC, (+)MiNP treatment/E-MC, and (+)MiNP/E-MC(FA). Fluorescent E-MC and E-MC(FA) uptake by 3 different cell subsets were quantified: non-immune cells (b, e), dendritic cells (c, f), and macrophages (d,g) for 4 different organs. Data are reported as fold increase median fluorescence intensity of E-MC or E-MC(FA) over a PBS baseline control. N=4-10 *p*<0.05. Significance was determined within each organ by separate unpaired student’s t-tests.

The administering of folate receptor targeted E-MC(FA) instead of E-MC had no significant effect when administered IV, but when administered SC, there was significantly increased uptake of E-MC(FA) in all tumor cell subsets (**Figure 4b-g**). This increased tumor accumulation for targeted nanoparticles following only one of the routes of administration was unexpected, but may be explained by differences in E-MC clearance in MPS organs. Only the SC injection of (+)MiNP resulted in significant decreases in macrophage and dendritic cell uptake of E-MC in the spleen and liver, which would account for the additional E-MC(FA) available for accumulate in tumors. Our in vitro data demonstrated that MiNP inhibits macropinocytosis but does not strongly impact receptor mediated endocytosis (Figure 2c,d, **Figure S5**), which we hypothesized would enhance the uptake of a receptor-targeted chase nanoparticle in vivo. Thus our co-administration of MiNP with E-MC possessing an additional folate receptor targeting element validated these in vitro results by significantly enhancing the accumulation of E-MC(FA) in B16F10 tumors by up to 8-fold (**Fig. 4 e-g**). This significant increase in uptake for E-MC(FA) versus E-MC was only observed within the solid tumors and in no other organs.

## Conclusion

Here, we demonstrate the nanoparticle biodistribution-altering effects of MiNP that encapsulate a small molecule inhibitor of macropinocytosis, LatA. We have characterized and evaluated these nanoparticles both in vitro and in vivo as key mediators in a co-administration strategy to increase the targeting efficacy of a second “effector” nanoparticle, E-MC. Clearance by the MPS remains a critical issue for many drug delivery applications beyond nanoparticles, suggesting a potentially broad range of applications for MiNP. For example, MPS organs are major sites of off-target accumulation for monoclonal antibodies^[33]^ and decreasing this effect may allow enhanced efficacy with lower dosages and fewer side effects during cancer therapy. Strategies for blocking the clearance of therapeutic antibodies have long been under investigation^[34]^, yet inhibiting non-specific uptake via macropinocytosis remains underexplored. Recently, the depletion of subcapsular sinus macrophages via liposomes loaded with clodronate and other agents was employed to investigate the role of these cells during nanovaccination^[35]^. Results showed that removal of these cells prior to immunization enhanced delivery of the nanovaccine to lymph node follicles for improved humoral responses. Our work supports such strategies while additionally demonstrating that inhibition of MPS cells can be performed in a reversible and nontoxic manner without killing phagocytes, many of which play critical downstream roles in the generation of an immune response. Furthermore, SC administration of MiNP increased the serum concentration of E-MC to levels similar to IV administration, potentially opening up new routes of administration and dosing regimens previously unavailable to many nanotherapeutics and controlled delivery strategies.

In a tumor model, we found that MiNP increase target tissue and cell accumulation through reduction of uptake by phagocytic cells such as macrophages and dendritic cells in the liver and spleen, which accounts for >90% of MPS cells. Our results validate LatA loaded PEG-*b*-PPS MiNP as a promising platform to improve the performance of other, paired effector nanoparticle therapeutic and diagnostic platforms. These proof-of-concept results justify the exploration of alternative MiNP formulations encapsulating inhibitors in addition to or in combination with LatA, as well as the investigation of MiNP as part of functional strategies employing effector nanoparticles loaded with diagnostic and/or therapeutic agents.

## Animal Experimentation

Mice were handled in accordance with protocol IS00001408 as approved by the Institutional Animal Care and Use Committee of Northwestern University.

## Experimental Section

Detailed experimental methods can be found in the supporting information.

## Acknowledgements

The authors are grateful to Jonathan Remis for cryo-TEM observation. We thank the support from the Center for Computation & Theory of Soft Materials, the BioCryo facility of Northwestern University’s NUANCE Center, the Integrated Molecular Structure Education and Research Center, Structural Biology Facility, NU Atomic, the Nanoscale Characterization Experimental Center, Robert H. Lurie Comprehensive Cancer Center Flow Cytometry Core, and Biological Imaging Facility at Northwestern University. SAXS experiments were performed at the DuPont-Northwestern-Dow Collaborative Access Team (DND-CAT) located at Sector 5 of the Advanced Photon Source (APS). DND-CAT is supported by Northwestern University, E.I. DuPont de Nemours & Co., and The Dow Chemical Company. This research used resources of the Advanced Photon Source, a U.S. Department of Energy (DOE) Office of Science User Facility operated for the DOE Office of Science by Argonne National Laboratory under Contract No. DE-AC02-06CH11357. Funding: This work was supported by the National Institutes of Health Director’s New Innovator Award (grant no. 1DP2HL132390-01), the National Science Foundation (CBET-1806007 and CAREER Award no. 1453576), and the Louis A. Simpson & Kimberly K. Querrey Center for Regenerative Nanomedicine Regenerative Nanomedicine Catalyst Award.

