## Supplemental data for "Co-administration of macropinocytosis inhibitory nanoparticles (MiNP) for enhanced nanoparticle circulation time and target tissue accumulation following subcutaneous injection"

| | Number Avg (nm) | PDI | LatA EE |
|---|---|---|---|
| (-)MiNP | 22.8±4.65 | 0.08 | N/A |
| (+)MiNP | 20.1±5.36 | 0.07 | 61.5% |

**Table S1.** Size and encapsulation efficiency (EE) of MiNP as determined by Dynamic Light Scattering (DLS) and HPLC. Data are presented for unloaded PEG-*b*-PPS micelles ((-)MiNP) and latrunculin A (LatA) loaded PEG-*b*-PPS micelles ((+)MiNP.

| | (-)MiNP | (+)MiNP |
|---|---|---|
| Radius of Core(nm) | 8.9 | 8.3 |
| Rg (nm) | 7.1 | 7.03 |
| Diameter | 32 | 30.6 |
| $Chi^2$ | 0.005 | 0.008 |

**Table S2.** Physical characteristics of MiNP as determined by small angle x-ray scattering (SAXS). Data are presented for unloaded ((-)MiNP) and latrunculin A loaded ((+)MiNP) micelles. All values including radius of gyration (Rg) were obtained using SAXS model fits.

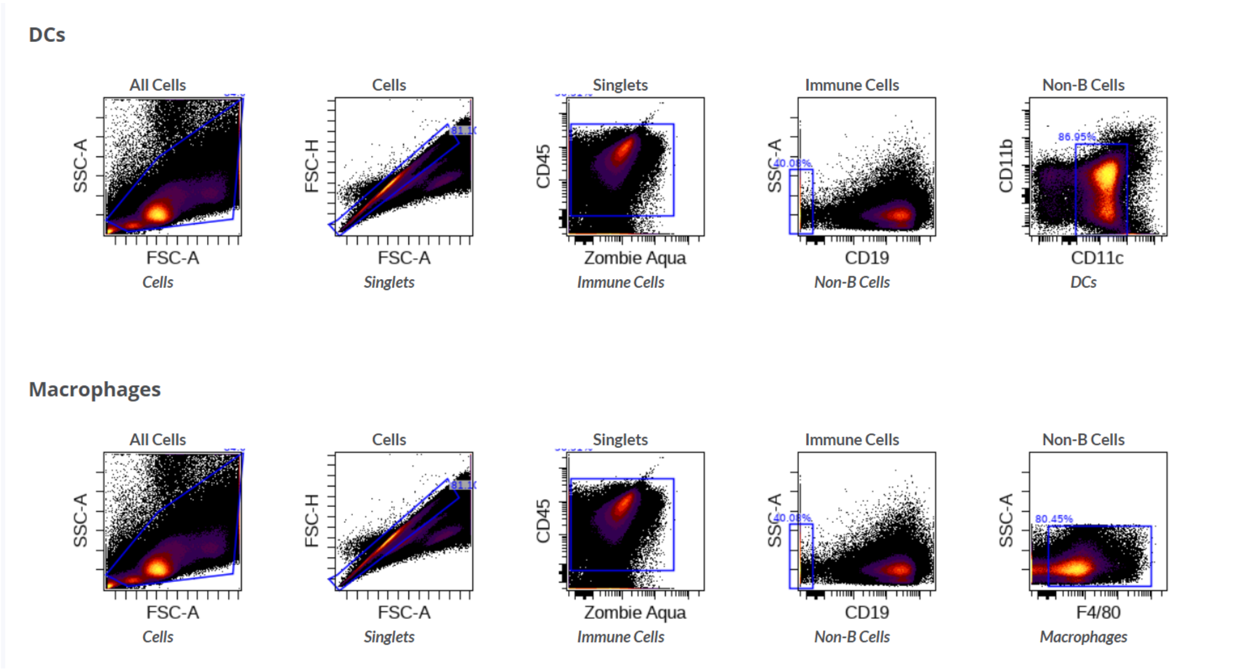


**Figure S1. Flow cytometry gating strategy for dendritic cells (DCs) and macrophages.** Cells were obtained from the spleen of mice injected with DiR labelled nanomaterials. Immune cells described in the manuscript were identified as follow: DCs: $CD45^+$, $CD11c^+$, $CD19^-$. Macrophages: $CD45^+$, $F4/80^+$, $CD19^-$.

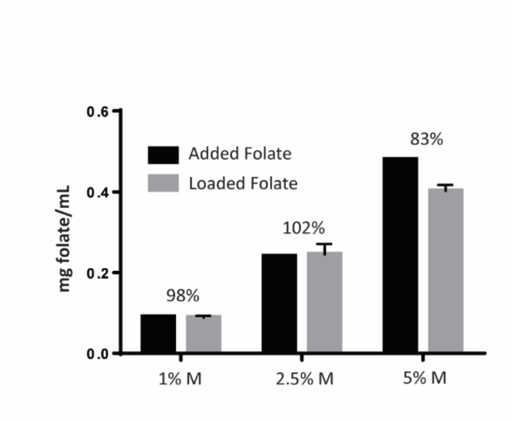


**Figure S2.** Custom lipid amphiphiles allow PEG-b-PPS micelles to incorporate and display folate with high efficiency. Folate-PEG-Lipid constructs were evaluated for their ability to partition into PEG-*b*-PPS micelles using spectroscopy by assessing the intrinsic absorption of folate. The amount of folate loaded onto micelles is shown in comparison to the amount of folate added for 3 different molar ratios of PEG-*b*-PPS polymer to folate-PEG-lipid construct**.**

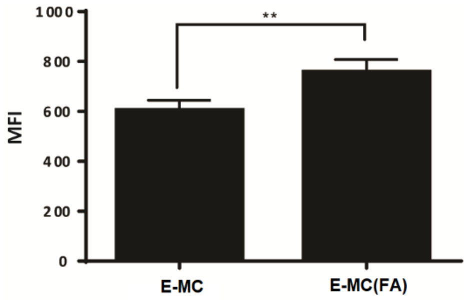


**Figure S3. Folate targeted ‘effector’ micelles (E-MC(FA)) are taken up at a higher rate by B16F10 cells than non-targeted E-MC.** B16F10 cells were incubated with non-targeted E-MC or folate receptor targeted E-MC(FA) for 1 hour and uptake was quantified by flow cytometry. Data are reported as median fluorescence intensity of E-MC or E-MC(FA). Statistical significance determined by unpaired student’s t-test. N=3, $p<0.005$.

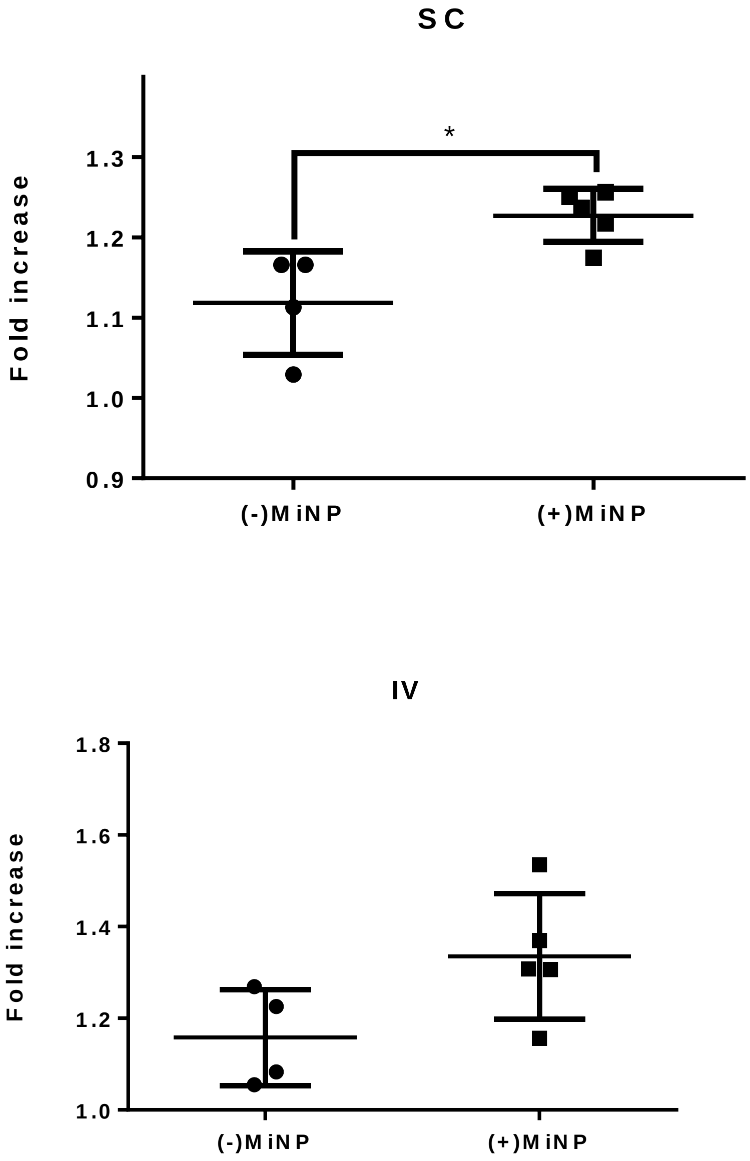


**Figure S4. Serum content of E-MC in tumor bearing mice after administration of either (+)MiNP or (-)MiNP 24 h post E-MC injection.** Data are expressed as fold increase of E-MC fluorescence over the baseline, which was defined by the average of 3 mice injected with nanoparticles lacking fluorescence. Significance was determined by unpaired student's t-test. N=4-5, $p<0.05$.

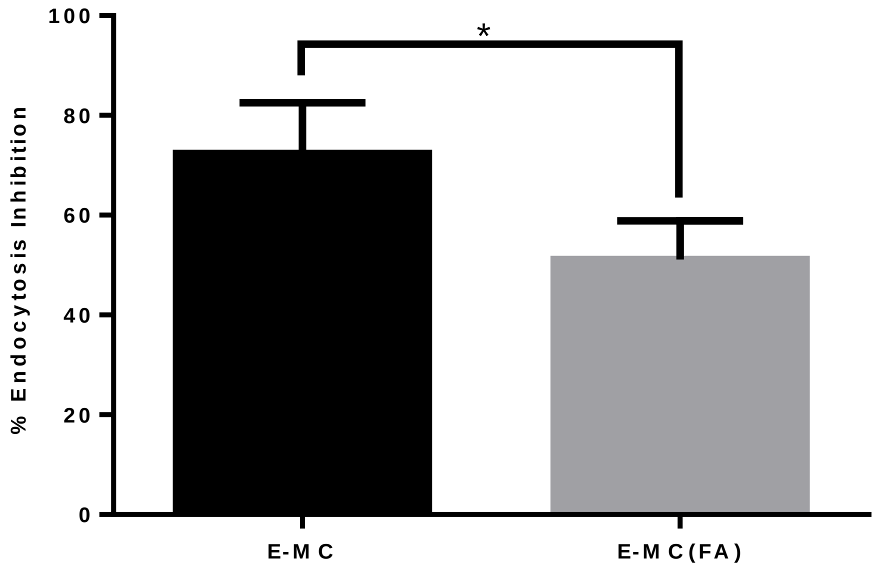


**Figure S5. Latrunculin A inhibits uptake of non-targeted E-MC more than targeted E MC(FA).** B16F10 melanoma cells were treated with 0.5 μM LatA for 2 h followed by 30 min of incubation with either non-targeted E-MC or folate receptor targeted E-MC(FA) prior to analysis by flow cytometry. Data are shown as percentage inhibition of endocytosis normalized to untreated macrophages. N=3 $p$<0.001

### Experimental Section

#### Synthesis of PEG-b-PPS copolymer

The self-assembling amphiphilic block copolymer $PEG_{45}$-b-$PPS_{23}$ was synthesized according to previously published protocols[1]. Briefly, PEG thioacetate was activated using sodium methoxide to initiate anionic ring opening polymerization of propylene sulfide. Following depletion of monomer, the propylene sulfide block was end capped with bromo benzene. Resulting polymer was double precipitated in methanol for purification and then evaluated using 1H NMR(CDCL3) and gel permeation chromatography (Thermo) using a waters styragel column with refractive index and UV-Vis detectors in a tetrahydrofuran (THF) mobile phase.

#### Synthesis of folate-PEG-palmitoleic acid targeting constructs

Folate targeting moieties were synthesized to selectively enhance the uptake of micelles by B16F10 melanoma cells. Folic acid-primary amine functionalized PEG (FA-PEG2k-NH2) was purchased from creative PEG works. Palmitoleic acid was covalently linked to FA-PEG2k-NH2 using an EDC reaction. Briefly, FA-PEG2k-NH2 was dissolved in DMF and EDC and palmitoleic acid were then added along with 4-dimethylamino pyradine as a catalyst. The reaction was allowed to run for 2 h and subsequently purified through double precipitation in cold diethyl ether. The resulting construct was characterized using 1H NMR (CDCL3) and gel permeation chromatography (Thermo) using a waters styragel column with refractive index and UV-Vis detectors in a THF mobile phase.

#### Assembly and loading of PEG-b-PPS micelles

Micelles were assembled from the previously described PEG-b-PPS block copolymer using a cosolvent evaporation method[1] .$PEG_{45}$-b-$PPS_{23}$ block copolymer, latrunculin A in various concentrations (Cayman Chemical) and 3uL DiI or DiR (Thermo)/10mg polymer(if fluorescence was required) were dissolved in 1mL dichloromethane (DCM). The resulting DCM solution was added dropwise to a scintillation vial containing 1 mL of sterile PBS stirred vigorously. DCM was allowed to evaporate over the course of 2 h. Micelles containing LatA were defined as (+)MiNP, while those without as (-)MiNP. Resulting micelles were then purified or set aside for addition of folate targeting moieties.

Folate targeted micelles (E-MC(FA)) were prepared through addition of 2.5% molar ratio of folate targeting moiety dissolved in DMSO. The micelle formulations were then placed on a shaker overnight to allow the palmitoleic acid tail to partition into the hydrophobic domain of the micelles. All formulations were then purified through gravity column chromatography using a Sephadex-LH20 column (Sigma) with a PBS mobile phase. Resulting micelles were then concentrated and further purified using a 10k MWCO Zebaspin desalting column (Thermo).

**Characterization of micelle formulations**

Once micelles were purified, their properties were characterized using a variety of techniques. Dynamic light scattering (DLS) was used to evaluate the zeta potential, size distribution, and polydispersity index (PDI) of the micelles using a zetasizer nano (Malvern Instrument) with a 4 mW He-Ne 633 nm Laser at 1 mg/mL in PBS. PDI was determined using a two-parameter fit to the DLS correlation data. Micelle morphology was characterized by cryogenic transmission electron microscopy (Gatan) (CryoTEM) as previously described[2]. Small angle X-ray scattering (SAXS) experiments were performed at the DuPont-Northwestern-Dow Collaborative Access

Team (DND-CAT) beamline at Argonne National Laboratory's Advanced Photon Source (Argonne, IL, USA). PRIMUS 2.8.2 software was utilized to obtain a final scattering curve after solvent buffer subtraction and structural parameters of micelle samples were determined using a polymer micelle model fit in SaxsView. LatA encapsulation and efficiency was determined using a previously described HPLC method[3]. 100 μL of concentrated and purified micelles were frozen at -80 °C and subsequently lyophilized overnight. Resulting cakes were solubilized in methanol to extract LatA. After 1 h of extraction, the suspension was centrifuged to eliminate precipitated polymer. Supernatant was then analyzed for LatA content using HPLC. A static Methanol:Water (95:5) mobile phase was used on an Agilent C18 XDB-Eclipse column with absorption (235nm) was used to evaluate LatA concentration. Standard curves were generated through serial dilution of LatA and subsequent lyophilization and methanol extraction of each standard.

Folate content of nanoparticles was determined using spectrophotometric analysis measuring absorption at 358 nm. Samples were analyzed in triplicate. Folate constructs in DMSO were serial diluted in PBS to obtain a standard curve. Blank micelles and PBS were used as baseline controls and their values were subtracted from FA-Micelles and FA-PEG-Palmitoleic acid standards respectively to create adjusted measurements to compensate for any interference from micelle polymer.

**Cell culture**

RAW 264.7 Macrophages were purchased from ATCC and cultured in T75 polystyrene tissue culture treated flasks (BD falcon) with DMEM (Life Technologies), 10% fetal bovine serum (FBS) (Gibco), and 1% penicillin/streptomycin (Life Technologies). Cells were passaged via mechanical cell scraping once they were 75-80% confluent.

B16F10 mouse melanoma cells were a generous gift from the laboratory of Dr. Bin Zhang at Northwestern University. B16F10 cells were cultured in T75 polystyrene tissue culture treated flasks (BD falcon) with DMEM (Life Technologies) plus 10% fetal bovine serum (FBS), and 1% penicillin/streptomycin (Life Technologies). Cells were passaged via trypsinization once 75-85% confluent. Both cell types were stored in an incubator at 37 °C and 5% CO2. Cells were passaged at a 1:4 ratio.

#### ***In Vitro* cell uptake assay**

Cell uptake assays were performed with both RAW 264.7 macrophages and B16F10 cells. Cells were adhered to 24 well tissue culture treated polystyrene plates (Costar) and treated with 0.25 mg of micelles containing DiI or DiR for 1 h at 37 °C and 5% $CO_2$. Cells were subsequently washed and harvested via mechanical scraping or trypsinization. Cells were then transferred to flow tubes and spun down at 300 x g for 5 min. Cells were subsequently stained with zombie aqua cell fixability dye (Biolegend) and fixed using IC fixation buffer (Biolegend). Flow cytometry was performed using FACS diva on a BD Fortessa flow cytometer (BD biosciences). Untreated cells of both types were used to subtract out background fluorescence, and flow data were analyzed using Cytobank software (Cytobank).

#### ***In vitro* macrophage cytotoxicity**

RAW264.7 macrophages were adhered to 24 well tissue culture treated polystyrene plates (Costar) and treated with free LatA or (+)MiNP at concentrations ranging from 0 μM to 10 μM. Cells were incubated at 37 °C and 5% $CO_2$. After 4 h, cells were washed and harvested via mechanical cell scraping. Cells were then stained with zombie aqua cell viability dye and fixed using IC fixation buffer (Biolegend). Flow cytometry was performed using FACS diva software on a BD Fortessa flow cytometer (BD biosciences). Ethanol killed RAW 264.7 macrophages stained with zombie aqua were used as a positive control. Flow data were analyzed using Cytobank Software (Cytobank).

**Cryogenic transmission electron microscopy**

Prior to applying sample, 200 mesh lacey copper grids were glow discharged. 4 µl of sample was applied to the grid. Blotting proceeded for 5 seconds with a blot offset of +0.5 mm, and was immediately plunged into liquid ethane using a FEI Vitrobot Mark III plunge freezing instrument. Grids were stored in liquid nitrogen. Samples were imaged using a JEOL JEM1230 LaB6 emission TEM operating at 100 keV. Plunge-frozen grids were held at -180 °C using a Gatan Cryo Transfer Holder. A Gatan Orius SC1000 CCD camera Model 831 was used to collect data. ImageJ software was used to process images.

**B16F10 melanoma tumor inoculation**

B16F10 melanoma cells were expanded using tissue culture treated T25 and T75 flasks. Once at an appropriate number, cells were harvested via trypsinization. Cells were counted and diluted to 3.5 million cells/mL and aliquoted in 500 µL volumes into sterile containers. C57/BL6 mice at 6 weeks of age were injected subcutaneously into the right hindquarters with 200 µL of the cell suspension. Mice were then returned to their enclosure and tumor growth was monitored over the next 10 days. Treatments were performed on days 12-14 after tumor inoculation.

**In vivo biodistribution studies**

All in vivo mouse studies followed one of two injection protocols. First, a pre-injection strategy was tested where (+/-)MiNP were administered twice at both 24 h and 4 h prior to injection of the chase effector nanoparticle. A simplified procedure was also evaluated wherein the (+/-)MiNP were administered alone just once to pre-condition the mouse, which was then followed 24 h later by a co-injected dose of a (+/-)MiNP and E-MC multi-nanoparticle formulation. Mice were then sacrificed at 24 h post (+/-)MiNP/E-MC injection. A dose of 100 μL of 7 μM (+)MiNP solution (approx. 0.55 mg/kg) was administered in each (+)MiNP injection. All injections were 100 μL in volume and the total injected amount of (+)MiNP or (-)MiNP and E-MC were equal.

Mice were sacrificed using cervical dislocation at 24 h and organs were harvested and placed in 24 well non tissue culture treated plates (Costar) in RPMI 1640 complete media (Gibco) on ice and were processed immediately. Once harvested, cells were isolated from each organ according to a different protocol (detailed below). Prior to staining, isolated cells were blocked with Cd16/32 blocking buffer for 15 min. Isolated cells were stained with CD11b (FITC), CD11c(BV421)), CD45(PerCp-Cy5.5), F4/80(PE-Texas Red), CD19 (Pe-Cy7) anti mouse antibodies, and zombie aqua cell viability dye (Amcyan) for 45 min. All cell staining materials were obtained from Biolegend. E-MC were labeled with Vybrant DiR (Thermo Fisher). Compensation controls were created from unstained mouse spleens. Cells were also isolated and stained from 3 mice that received no treatment. These mice were used as a baseline fluorescence control and used to calculate the relative fluorescence increase of E-MC in each of the different cell subsets (fold increase in Figure 3 and Figure 4). After staining, cells were washed and fixed with 1:1 cell staining buffer and cell fixation buffer and were processed using flow cytometry within 3 days of fixation. Flow cytometry data was analyzed using Cytobank software (Cytobank) and the gating strategy is shown in figure S1. Data are reported as fold increase of E-MC median fluorescence intensity over median fluorescence intensity of untreated mice. Significance was determined using students t-test within each cell subset.

**Liver**

Livers were incubated at 37 $^{\circ}$C in 0.2 mg/mL Dnase (Sigma), 5000 U/mL Collagenase IV (Sigma) solution in RPMI media (Gibco) for 1 h with agitation every 10 min. After chemical digestion, livers were mechanically disrupted through a 40 micron cell strainer and the resulting suspensions were centrifuged at 50 x g for 5 minutes. The supernatant was reclaimed and centrifuged at 50 x g a second time and the supernatant was again reclaimed. The resulting suspension was then centrifuged at 400 x g for 5 min to pellet remaining cells at which point the pellet was resuspended in red blood cell lysis buffer (Biolegend) and allowed to rest at 4 $^{\circ}$C for 15 min. Then the suspension was diluted with PBS and centrifuged at 400 x g to pellet the cells. Cells were then resuspended in cell staining buffer and transferred to flow tubes for staining.

**Spleen**

Spleens were mechanically disrupted and passed through a 40 micron cell strainer with RPMI media (Gibco). The resulting cell suspension was centrifuged at 400 x g to pellet cells and then resuspended in RBC lysis buffer (Biolegend). This suspension was incubated for 15 min at 4 $^{\circ}$C, then diluted with PBS and centrifuged at 400 x g for 5 min. The resulting pellet was then resuspended in cell staining buffer and transferred to flow tubes for staining.

**Lymph nodes**

Draining lymph nodes were mechanically disrupted and passed through a 40 micron cell strainer with RPMI media (Gibco). The resulting cell suspension was centrifuged at 400 x g to pellet cells. The pellet was then resuspended in cell staining buffer and transferred to flow tubes for staining.

**B16F10 tumor**

Tumors were diced using biopsy punches and the resulting tissue sections were incubated at 37°C in 0.2 mg/mL Dnase (Sigma), 5000 U/mL Collagenase IV (Sigma) solution in RPMI media (Gibco) for 1.5 h with agitation every 10 min. After chemical digestion, tumor tissue and collagenase solution were passed through a 40 micron cell strainer and the resulting cell suspensions were centrifuged at 50 x g for 5 min. The supernatant was reclaimed and centrifuged at 50 x g a second time and the supernatant was again reclaimed. The resulting cell suspension was then centrifuged at 400 x g for 5 min to pellet remaining cells at which point the pellet was resuspended in red blood cell lysis buffer (Biolegend) and allowed to rest at 4 ºC for 15 min. The resulting suspension was diluted with PBS and centrifuged at 400 x g to pellet the cells. Cells were then resuspended in cell staining buffer and transferred to flow tubes for staining.

**Serum**

Whole blood was obtained from mice via retroorbital bleed. Approximately 100 μL of blood was obtained at each bleed. Immediately after whole blood was collected, it was allowed to coagulate at room temperature for 30 min. Samples were then centrifuged for 10 min at

1500 x g in a centrifuge at 4ºC. Supernatant was isolated immediately after centrifugation and stored at 4ºC until analysis, which was no more than 4 h after serum isolation.

**Statistical analysis**.

All statistical analyses were performed using GraphPad Prism software (GraphPad Software; version 8.4.1). Each figure notes the specific type of statistical test utilized to determine statistical significance.

### Author Contributions

All experiments were designed by TS and EAS. PEG-*b*-PPS polymer was synthesized by MF. Micelle formulations were made by TS. TEM and DLS was performed by MV. SAXS data was collected and analysed by SB. Animal work including injections, care, and endpoint assays were performed by YL, TS, and SB. Flow cytometry, antibody panel design and data analysis were done by TS. In vitro uptake and cytotoxicity assays were performed by TS. Figures were designed by TS, MV, MF, and EAS. Manuscript was written by TS, SB, and EAS.
